# Cas4/1 dual nuclease activities enable prespacer maturation and directional integration in a type I-G CRISPR-Cas system

**DOI:** 10.1101/2023.06.05.543779

**Authors:** Yukti Dhingra, Dipali G. Sashital

## Abstract

CRISPR-Cas adaptive immune systems uptake short ‘spacer’ sequences from foreign DNA and incorporate them into the host genome to serve as templates for crRNAs that guide interference against future infections. Adaptation in CRISPR systems is mediated by Cas1-Cas2 complexes that catalyze integration of prespacer substrates into the CRISPR array. Many DNA targeting systems also require Cas4 endonucleases for functional spacer acquisition. Cas4 selects prespacers containing a protospacer adjacent motif (PAM) and removes the PAM prior to integration, both of which are required to ensure host immunization. Cas1 has also been shown to function as a nuclease in some systems, but a role for this nuclease activity in adaptation has not been demonstrated. We identified a type I-G Cas4/1 fusion with a nucleolytically active Cas1 domain that can directly participate in prespacer processing. The Cas1 domain is both an integrase and a sequence-independent nuclease that cleaves the non-PAM end of a prespacer, generating optimal overhang lengths that enable integration at the leader side. The Cas4 domain sequence-specifically cleaves the PAM end of the prespacer, ensuring integration of the PAM end at the spacer side. The two domains have varying metal ion requirements. While Cas4 activity is Mn^2+^ dependent, Cas1 preferentially uses Mg^2+^ over Mn^2+^. The dual nuclease activity of Cas4/1 eliminates the need for additional factors in prespacer processing, making the adaptation module self-reliant for prespacer maturation and directional integration.

## Introduction

CRISPR-Cas (clustered regularly interspaced short palindromic repeats, CRISPR-associated) systems are adaptive immune systems that immunize prokaryotes against mobile genetic elements by acquiring short DNA sequences from invaders as “spacers” in CRISPR arrays (1–3). CRISPR arrays comprise a leader sequence followed by a series of repeats interspersed with spacers (4). New spacers are inserted at the junction of the leader and the first repeat, allowing for chronological spacer insertion (4–7). Transcription of the CRISPR and subsequent processing generates CRISPR RNAs (crRNAs) that guide Cas effectors for target recognition and degradation (8, 9). Thus, new spacer acquisition underlies downstream expression and interference ensuring a robust immune response and neutralization of future infections (10, 11).

All CRISPR-Cas systems that perform spacer acquisition contain Cas1 and Cas2 proteins that form an integration complex (4, 6, 12–15). Several DNA-targeting type I, II and V CRISPR-Cas systems also contain Cas4 (16), an accessory adaptation protein that can either associate with the Cas1-Cas2 complex, forming a Cas4-Cas1-Cas2 complex (17), or can be fused to Cas1 (18), allowing formation of a Cas4/1-Cas2 complex (19). In these systems, Cas4 is required for acquiring spacers that can be subsequently used for DNA targeting by a Cas effector (20–24). DNA-targeting Cas effectors recognize short 3-5 base pair protospacer adjacent motif (PAM) sequences to rapidly scan through invading foreign DNA and locate the crRNA-complementary target, a process that is required for eventual target binding and degradation (25–29). Thus, spacers acquired in DNA-targeting CRISPR-Cas systems must be acquired from regions that contain a PAM to encode a crRNA that can successfully immunize the host (30). Synthesis of a crRNA complementary to the target also requires that the PAM is precisely removed from the prespacer and that the prespacer be inserted in the proper orientation at the leader-repeat junction (10, 11).

Cas4 allows the adaptation complex to achieve PAM-specific prespacer selection and processing (17, 22–24, 31), along with directional integration (19, 21, 32, 33). Cas4 is a PAM-specific endonuclease that recognizes PAM sequences in prespacers within its active site, but delays cleavage just upstream of the PAM until integration is initiated (17, 19, 32). Cas1 catalyzes two transesterification reactions to mediate insertion of the 3′ ends of the prespacer substrate into the CRISPR array, at the leader repeat junction (leader side) and the repeat spacer junction (spacer side) (5). Integration of the non-PAM end of the prespacer at the leader side precedes PAM cleavage by Cas4 and PAM end integration at the spacer side by Cas1 (19). Cas4 remains bound to the PAM end of the prespacer following leader side integration, ensuring that the prespacer is integrated in the proper orientation (19, 32).

Processing of the non-PAM end of the prespacer is also necessary to achieve an optimal overhang length for integration by Cas1. Cas1-Cas2 optimally integrates prespacers with 22-23 base pair (bp) duplexes and 5-7 nucleotide (nt) 3′ overhangs (6, 13, 34). Host exonucleases like DnaQ and ExoT have been shown to trim prespacer overhangs (35, 36). In some Cas4-less systems, DnaQ is fused to the Cas2 subunit (37–39), enabling prespacer trimming by subunits of the adaptation complex (37). Cas1 can also act as a metal dependent sequence nonspecific endonuclease in some systems, suggesting that Cas1 could also be involved in prespacer processing (34, 40, 41). However, prespacer processing by Cas1 in a Cas4-containing system has not been reported.

Here, we investigated the nuclease activities of both the Cas1 and Cas4 domains of a Cas4/1 fusion the *Methanosarcina barkeri* type I-G system. We demonstrate that the Cas1 domain can function as a Mg^2+^-dependent sequence nonspecific nuclease and process DNA overhangs, producing prespacers that are optimal for integration. In contrast, Cas4 cleaves the PAM-containing overhangs using a sequence-specific and Mn^2+^-dependent mechanism to ensure specific prespacer selection and orientation of integration. This dual nuclease activity of Cas4/1 enables prespacer maturation and directional integration that is self-contained within the adaptation module.

## Results

### Cas4/1 fusion has dual nuclease activities with varying metal dependence

To investigate the nuclease activities of a Cas4/1 fusion, we recombinantly expressed and purified Cas4/1 and Cas2 from a type I-G system in *Methanosarcina barkeri str. Weismoor* (hereafter MbCas4/1 and MbCas2, **Fig. 1A**). We tested the ability of Cas4/1 to process prespacer substrates using a panel of metal-ion cofactors that are commonly found to support nuclease activities (**Fig. S1A-B**). The prespacer contained a 22 bp duplex and 15 nucleotide (nt) 3′ overhangs. One strand of this prespacer contained a 5′ - GAA - 3′ PAM beginning at the eighth position of the overhang (PAM strand) and the other strand lacked a PAM in the overhang (non-PAM strand) (**Fig. 1B, S1A**). In the presence of MbCas4/1, we observed cleavage products with all metal ions except Zn^2+^ (**Fig. S1B**). Cleavage occurred in both the absence and presence of Cas2 for most metal ions, with the exception of Co^2+^, where cleavage was much weaker in the absence of Cas2.

**Figure 1.**
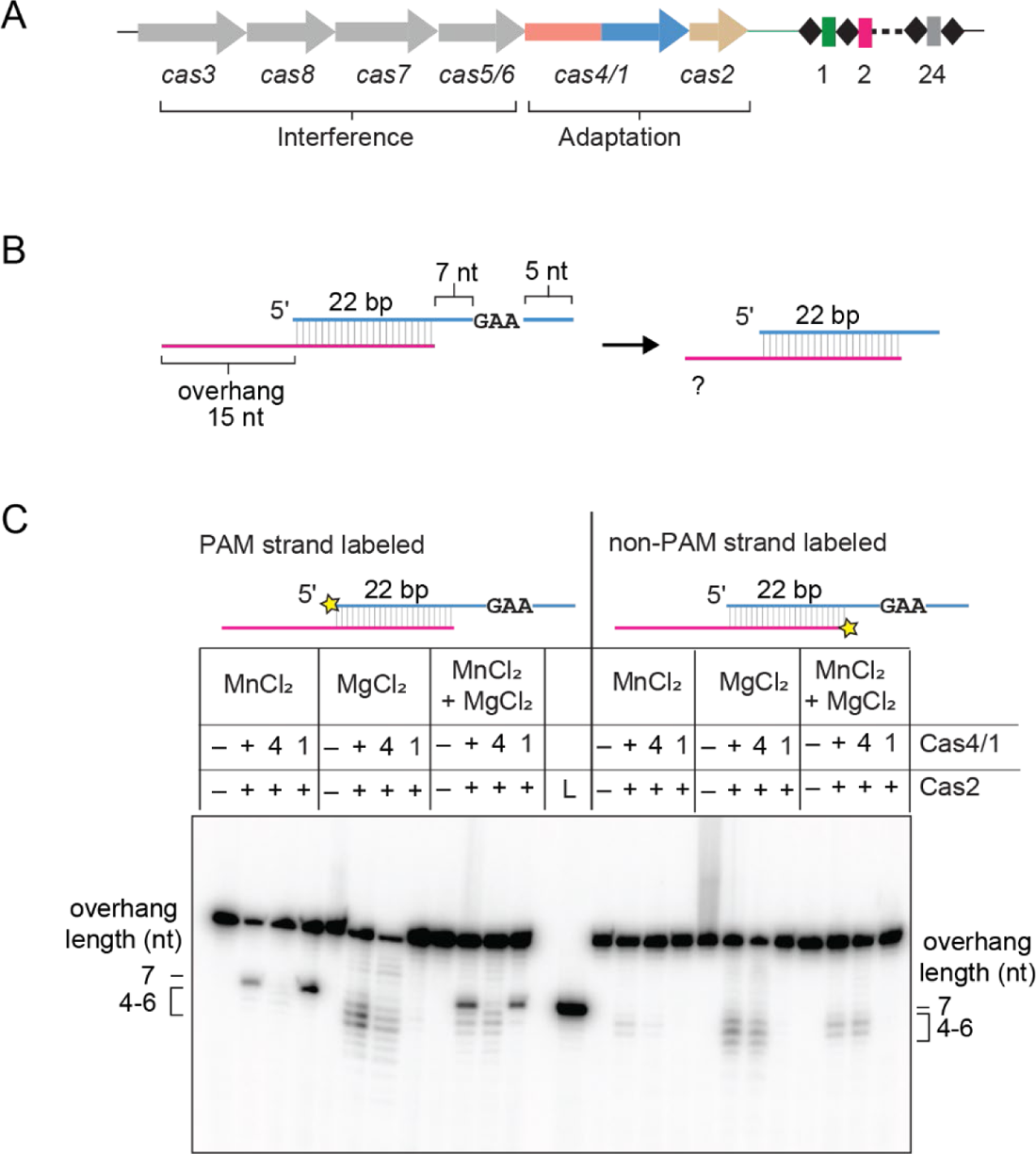
Cas4/1 fusion has dual nuclease activity with varying metal dependence. **A,** Schematic of *cas* genes and CRISPR locus of type I-G system in *Methanosarcina barkeri str. Weismoor. cas* genes are shown as colored arrows with the gene products involved in adaptation or interference indicated. Spacers and repeats in the CRISPR array are shown as colored rectangles and black diamonds, respectively. **B,** Schematic of prespacer cleavage assay for a PAM/NoPAM substrate with a 5′-GAA-3′ PAM sequence. **C.** Denaturing polyacrylamide gel showing cleavage assay with substrate shown in (B) with PAM or non-PAM strand labeled. The radioactive label is shown with a yellow star. Cas4 domain (E101A) and Cas1 domain (E375A) active site mutants are indicated as 4 and 1, respectively. Overhang lengths for cleavage products are indicated. A control DNA corresponding to the expected Cas4 cleavage products was run in the lane labeled L. Metal ion concentrations used for the three conditions are 2 mM MnCl_2_; 2 mM MgCl_2_; 1 mM MgCl_2_ and 0.5 mM MnCl_2_.

The nuclease activity of the type I-G Cas4/1 fusion protein from *Geobacter sulfurreducens* (hereafter GsCas4/1) has previously been shown to be strictly Mn^2+^ dependent (19). The more promiscuous metal dependence of MbCas4/1 suggests either that the Cas4 domain can tolerate other metal ions, or that another nuclease activity is present in the protein, potentially within the Cas1 domain. To test this, we introduced substitutions to active site residues in the Cas4 (E101A) and Cas1 (E375A) domains of MbCas4/1. We tested the cleavage activities of wild-type (WT) Cas4/1 and the two active site mutants in the presence of Mn^2+^, Mg^2+^ or both (**Fig. 1C**). We used a substrate containing a PAM and a non-PAM strand, in which one strand was radioactively labeled (**Fig. 1B, C**).

When the PAM strand was labeled, we observed a prominent cleavage product in the presence of Mn^2+^ that was consistent with the size of the expected Cas4 product upon cleavage just upstream of the PAM, yielding a 7 nt overhang (**Fig. 1B, C**). A similarly sized product was not observed when the Cas4 active site mutant was used, when the non-PAM strand was labeled, nor when only Mg^2+^ was present in the reaction. These results are consistent with previous studies of GsCas4/1, suggesting that the Cas4 domain of MbCas4/1 is also a Mn^2+^-dependent, PAM-specific endonuclease (19). In the presence of Mg^2+^, we observed multiple cleavage products corresponding to overhang lengths of 4-7 nt for both the PAM and non-PAM strands with both the wild-type and Cas4-domain active site mutant (**Fig. 1C**). These cleavage products were not observed with the Cas1-domain active site mutant. Overall, these results indicate that the Mg^2+^-dependent nuclease activity lies within the Cas1 domain and that this activity is not dependent on the presence of a PAM within the prespacer overhang.

To determine whether both Cas1 and Cas4 nuclease domain activities could be observed at physiologically relevant metal ion conditions, we titrated the concentration of Mg^2+^ with a constant concentration of Mn^2+^ and vice versa (**Fig. S1C-D**). We could readily observe both nuclease activities at 2 mM Mg^2+^ and 0.2 mM Mn^2+^. Cas4-dependent cleavage was less pronounced at 0.1 mM Mn^2+^, although a small amount of Cas4-dependent product could still be observed when the Cas1 domain mutant was used. Cas1 can use both Mn^2+^ and Mg^2+^ for its activity, however we observed reduced activity of Cas1 above 0.5 mM Mn^2+^. Our results strongly suggest that MbCas4/1 exhibits dual nuclease activity where the Cas4 domain requires Mn^2+^ but the Cas1 domain preferentially uses Mg^2+^ over Mn^2+^ for its nucleolytic activity.

### Cas1 nuclease activity is sequence independent

We previously showed that Cas4 from the *Alkalibacillus halodurans* (hereafter AhCas4) type I-C system can process prespacer substrates where the PAM is located at different locations within the overhang (17). PAM-specific cleavage results in products of different lengths, as AhCas4 always cuts just upstream of the PAM sequence. Using a similar set of substrates (**Fig. 2A**), we tested the activity of MbCas4/1 in the presence of Mn^2+^ or Mg^2+^ to understand the sequence specificities of the two nuclease activities. We designed four 22 bp duplex substrates containing a non-PAM strand with a 7-nt overhang and a PAM strand with 3-9 nt of single-stranded DNA between the duplex and the PAM (**Fig. 2A**). In the presence of Mn^2+^, Cas4-mediated cleavage products for the four different substrates varied in size depending on the location of PAM, consistent with our previous observations (**Fig. 2B**). On the other hand, cleavage products resulting from Cas1 activity in the presence of Mg^2+^ were similarly sized, ranging from overhangs of 4-7 nt for all substrates. This suggests that while the Cas4-mediated endonuclease activity is PAM-dependent, as in other Cas4 and Cas4/1 proteins, the Cas1-mediated nuclease activity is sequence independent and instead length dependent.

**Figure 2.**
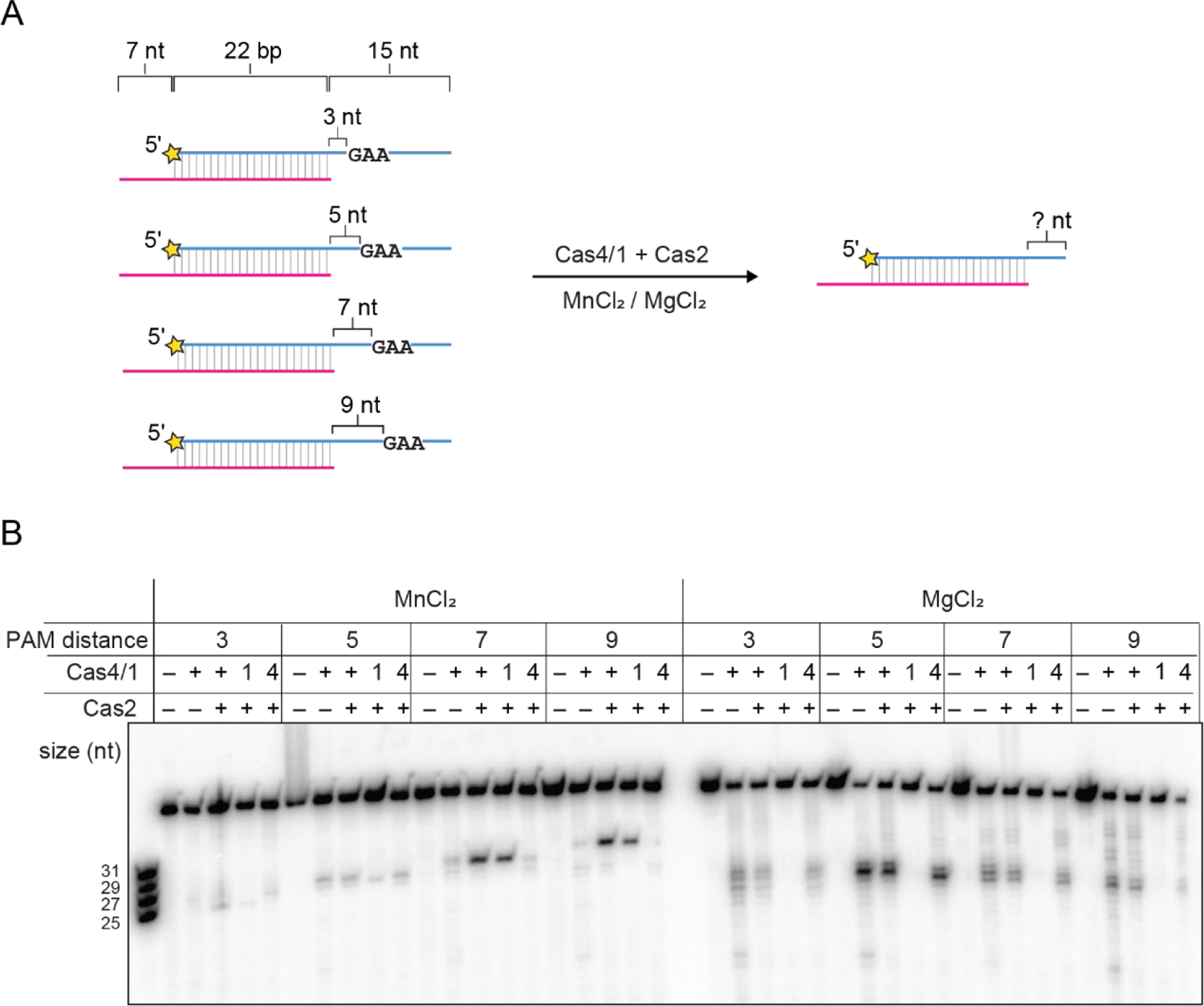
Cas4 nuclease activity is sequence dependent and Cas1 nuclease activity is sequence independent. **A,** Schematic of cleavage assay with Cas4/1 and Cas2 in the presence of 2 mM MnCl_2_ or 2 mM MgCl_2_ for a panel of substrates with variable distance between the duplex and PAM. The 5′-GAA-3′ PAM is located 3, 5, 7, or 9 nt after the end of the duplex. Radioactive label is indicated with a yellow star. **B,** Denaturing polyacrylamide gel showing cleavage assay with substrates shown in (A). Cas4 domain (E101A) and Cas1 domain (E375A) active site mutants are indicated as 4 and 1, respectively. The ladder on the left includes the four expected Cas4 cleavage products for the four different substrates.

### Cas4 cleavage is independent of leader side integration and enables spacer side integration

For a spacer to be functional, the non-PAM end of the prespacer must be integrated at the leader side and the PAM end must be integrated at the spacer side. It has previously been shown that Cas4 remains bound to the PAM end of the prespacer following leader side integration in both type I-G and I-C systems (19, 32). For type I-G GsCas4/1, the Cas4 domain is activated for PAM cleavage only after leader side integration of the non-PAM end (19), although a similar activation was not observed for type I-C AhCas4 (32). These experiments were performed using a half-site integration (HSI) intermediate where one strand of the prespacer is already integrated at the leader side and the other strand contains an overhang with a PAM (**Fig. 3A**). For GsCas4/1, use of such a substrate significantly enhanced Cas4-dependent PAM processing compared to an unintegrated prespacer substrate (19).

**Figure 3.**
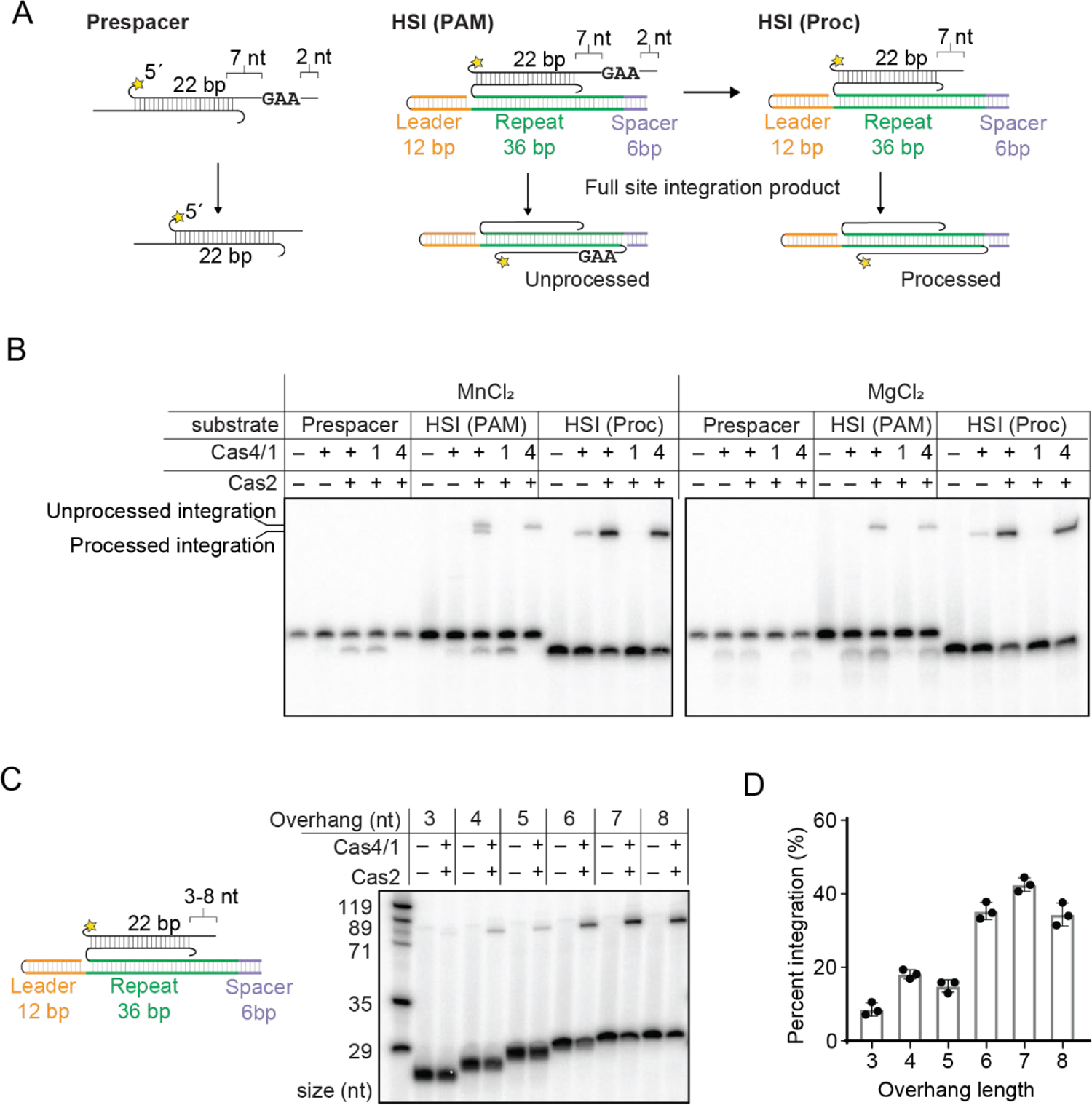
Cas4 cleavage is independent of leader side integration and creates optimal substrates for spacer side integration. **A,** Schematic of substrate and product for a cleavage and integration assay with half-site intermediates (HSI) of a prespacer containing an unprocessed PAM strand or a pre-processed strand. Radiolabel is indicated with a star in the substrate design. **B,** Denaturing polyacrylamide gel showing cleavage/integration assay with substrates shown in (A) in the presence of 2 mM MnCl_2_ or 2 mM MgCl_2_. Cas4 domain (E101A) and Cas1 domain (E375A) active site mutants are indicated as 4 and 1, respectively. Integration products for unprocessed and processed substrates are indicated. **C,** Schematic of substrate with variable length of overhang used for integration assay. Radiolabel is indicated with a yellow star. 2 mM MgCl_2_ was used for this assay. Denaturing polyacrylamide gel showing spacer side integration assay with substrates shown on left. Sizes of ladder are indicated. **D,** Quantification of integration assay shown in (C). Data is representative of three independent replicates. Error bars represent standard deviation, and the three data points are plotted as dots.

To determine if MbCas4/1 uses a mechanism of Cas4 cleavage activation upon leader side integration, we performed a similar experiment, comparing the cleavage activities of both Cas1 and Cas4 domains in MbCas4/1 with an unintegrated prespacer or the HSI substrate containing a PAM (**Fig. 3A-B**). We observed similar cleavage activities for both the prespacer substrate and the HSI substrate containing a PAM in the presence of both Mn^2+^ and Mg^2+^. This suggests that the Cas4 domain of MbCas4/1 is not activated solely by half-site integration, as was observed for GsCas4/1 (19).

The HSI substrate also enables observation of spacer-side integration based on the formation of slower migrating DNA species representing the full-site integration (FSI) product. We observed two FSI products in the presence of Mn^2+^ when using WT Cas4/1 (**Fig. 3B**). The shorter product corresponds to the length of the FSI product produced using a pre-processed HSI substrate, while the longer product corresponds to the length of the FSI product produced when Cas4 activity is ablated. Thus, these two products are consistent with spacer side integration of both the unprocessed and processed strand.

In contrast, in the presence of Mg^2+^, we observed integration of unprocessed prespacer but could not detect shorter integration products, despite observing Cas1 cleavage products (**Fig. 3B**). The Cas1 cleavage products ran as more diffuse and faster migrating bands than the Cas4 product, suggesting that Cas1 cleavage of the HSI substrate produces overhangs of multiple lengths that are shorter than 7 nt, similar to the products observed for the prespacer substrate (**Fig. 1C**). This raises the possibility that the Cas1 cleavage products might be shorter than the optimal length required for spacer-side integration. To determine the optimal length of overhangs for Cas1-mediated integration, we designed a panel of HSI substrates with variable overhang lengths of the processed strand between 3 to 8 nt (**Fig. 3C-D**). We observed a higher percentage of spacer-side integration for overhang lengths of 6 to 8 nt and very low percent integration for overhangs of 3 to 5 nt. These results suggest that Cas4-mediated cleavage products can be integrated at the spacer side as long as the PAM is 7 nt or further away from the duplex, enabling integration of the properly processed PAM strand. However, Cas1-mediated cleavage of overhangs in HSI substrates may produce products that are too short to be optimally integrated at the spacer side, preventing integration of PAM strands that are inadvertently processed by Cas1.

### Cas4/1 enables directional integration

Our results suggest that Cas4 is required to create optimal overhangs for spacer-side integration, but do not rule out the possibility that Cas1 cleavage of the non-PAM overhang could enable leader-side integration. Thus, processing of the two ends of the prespacer could be achieved by the two different Cas4/1 nuclease activities. Cas1 cleavage and integration of the non-PAM end must occur prior to Cas4 cleavage of the PAM end to ensure directional integration.

To test this model, we used a plasmid integration assay. In this assay, prespacers are integrated into a plasmid carrying the full leader and one repeat for the type I-G CRISPR in *Methanosarcina barkeri* (**Fig. 4A**). We assessed integration of prespacers at either site using PCR and high throughput sequencing analysis of the resulting amplicons. For the PCR, we used primers specific to one of the two prespacer strands and either the leader (spacer side integration S1 and S2) or the plasmid backbone (leader side integration L1 and L2). All four combinations of primers (L1, L2, S1 and S2) yield a 175 bp product upon integration at the correct sites in the CRISPR (**Fig. 4B**). Integration of the non-PAM strand at the leader side (detected using the L2 primer combination) and the PAM strand at the spacer side (detected using the S1 primer combination) indicates a correctly oriented prespacer.

**Figure 4.**
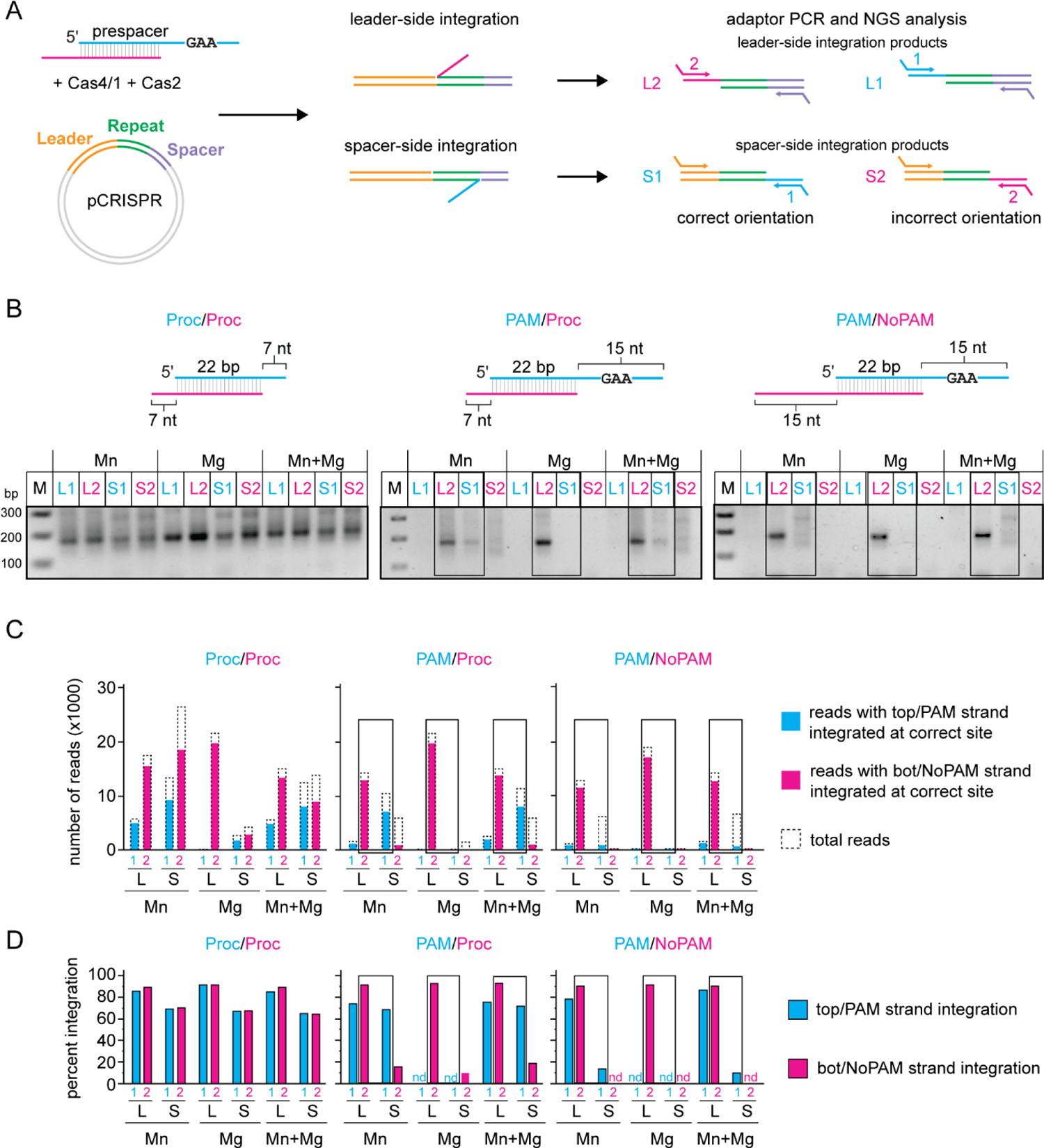
Cas4/1 achieves directional integration of prespacers. **A,** Schematic of plasmid integration assay. pCRISPR contains the leader sequence and one repeat sequence from the type I-G CRISPR in *M. barkeri.* The two integration events, leader side and spacer side, are shown. Only integration of one strand is shown at a time for simplicity. Four possible combinations of products are shown for top/PAM strand (cyan) and bottom/NoPAM strand (magenta) integrated at leader side or spacer side. Arrows represent primers for PCR reactions with adaptor overhangs. L1 and L2 represent integration of top and bottom strand at leader side, respectively. S1 and S2 represent integration of top or bottom strand at spacer side, respectively. L2 and S1 products are expected for integration in the correct orientation. **B,** PCR products for integration reactions for Proc/Proc (both ends pre-processed), PAM/Proc (unprocessed PAM end and pre-processed non-PAM end) and PAM/NoPAM (unprocessed PAM and non-PAM end) substrates into pCRISPR in the presence of only 2 mM Mn^2+^, only 2 mM Mg^2+^, or both 1 mM MgCl_2_ and 0.5 mM MnCl. **C and D,** Illumina MiSeq data analysis for PCR products shown in (B). (C) Number of total reads (dotted line) and reads for correctly integrated top (cyan bar) or bottom (magenta bar) strand plotted for all four combinations of integration events (L1, L2, S1 and S2). (D) Percent of integration events at the correct leader or spacer side site for all four combinations of integration events. “nd” indicates that < 40 total reads were detected for the sample. Black boxes in (B), (C), and (D) indicate integration events representative of correct orientation of integration.

We tested integration of three prespacers in the presence of only 2 mM Mn^2+^, only 2 mM Mg^2+^, or both 1 mM Mg^2+^ and 0.5 mM Mn^2+^ (**Fig. 4B**). As expected and consistent with previous results on Cas1-Cas2 mediated integration (5, 35), we observed PCR products for integration of either strand at the leader and spacer side for a prespacer in which both strands contained 7 nt processed overhangs (Proc/Proc). For prespacers containing a PAM strand, we observed a PCR product with the L2, but not the L1, primer combination for all three metal conditions. These results suggest that leader side integration occurs only with the non-PAM strand of the prespacer, but not the PAM strand (**Fig. 4A**). Conversely, we only observed a 175 bp PCR product for the substrate with a pre-processed non-PAM end (PAM/Proc) using the S1, but not the S2, primer combination, and only when Mn^2+^ was present during integration (**Fig. 4B**). These results are consistent with PAM end integration at the spacer side, and most likely occurring only after cleavage by Cas4 in a Mn^2+^-dependent manner. For the S2 combination, as well as for the S1 combination using a prespacer with an unprocessed non-PAM end (PAM/NoPAM), a smear of PCR products appeared on the gel, suggesting that integration may have occurred at nonspecific locations.

We next subjected the PCR amplicons to high-throughput sequencing (**Fig. 4A**). We first analyzed the frequency of integration for both PAM and non-PAM strands at the correct site on both the leader and spacer sides (**Fig. 4C, D**). Figure 4C plots the number of reads with the specific integration events, while Figure 4D plots the same values as percentage of total reads. We observed unoriented integration of a fully processed prespacer (Proc/Proc), consistent with the PCR results (**Fig. 4B-D**). Similar to other systems (7, 22, 42), integration at the leader side was more specific than integration at the spacer side, with 80-90% of integration events occurring at the correct leader side site and only 60-70% of integration events occurring at the correct spacer side site (**Fig. 4D**). We similarly observed high-frequency and high-fidelity leader side integration for non-PAM strands of PAM-containing substrates (PAM/Proc and PAM/NoPAM, **Fig. 4C, D**). In contrast, leader side integration of the PAM strand occurred very rarely and only in the presence of Mn^2+^ (**Fig. 4C**), and was lower fidelity (∼74% integration at the correct site) than for the non-PAM strand (**Fig. 4D**). Instead, PAM end integration mainly occurred at the spacer side in the presence of Mn^2+^ (**Fig. 4C**), with ∼70% of integration events occurring at the correct site for the prespacer containing a processed non-PAM end (PAM/Proc) (**Fig. 4D**). However, for the prespacer containing an unprocessed non-PAM end (PAM/NoPAM), integration of the PAM end only occurred at the correct site for 10-13% of reads in the presence of Mn^2+^. Notably, spacer side integration of the non-PAM end of the PAM/NoPAM substrate was not detected (**Fig. 4C**), indicating that only the PAM end of this substrate was integrated at the spacer side, albeit inefficiently. We did not detect integration of the PAM end of either substrate when only Mg^2+^ was present, indicating that cleavage by Cas4 is essential for integration of the PAM end (**Fig. 4C, D**). Overall, these results suggest that the Cas4 domain prevents premature integration of the PAM end at the leader side, enabling directional integration of prespacers into the CRISPR array.

### Cas4/1 dual nuclease activities process both ends of the prespacer prior to integration

We next investigated whether prespacers were processed by either Cas4 or Cas1 prior to integration. For the L2 and S1 primer pair PCR products, we determined the overhang length of prespacers that were integrated at the correct site (**Fig. 5A**). The number of reads were plotted against the overhang length for the non-PAM strand and PAM strand integrated at the leader and spacer sides, respectively (**Fig. 5B-G**).

**Figure 5.**
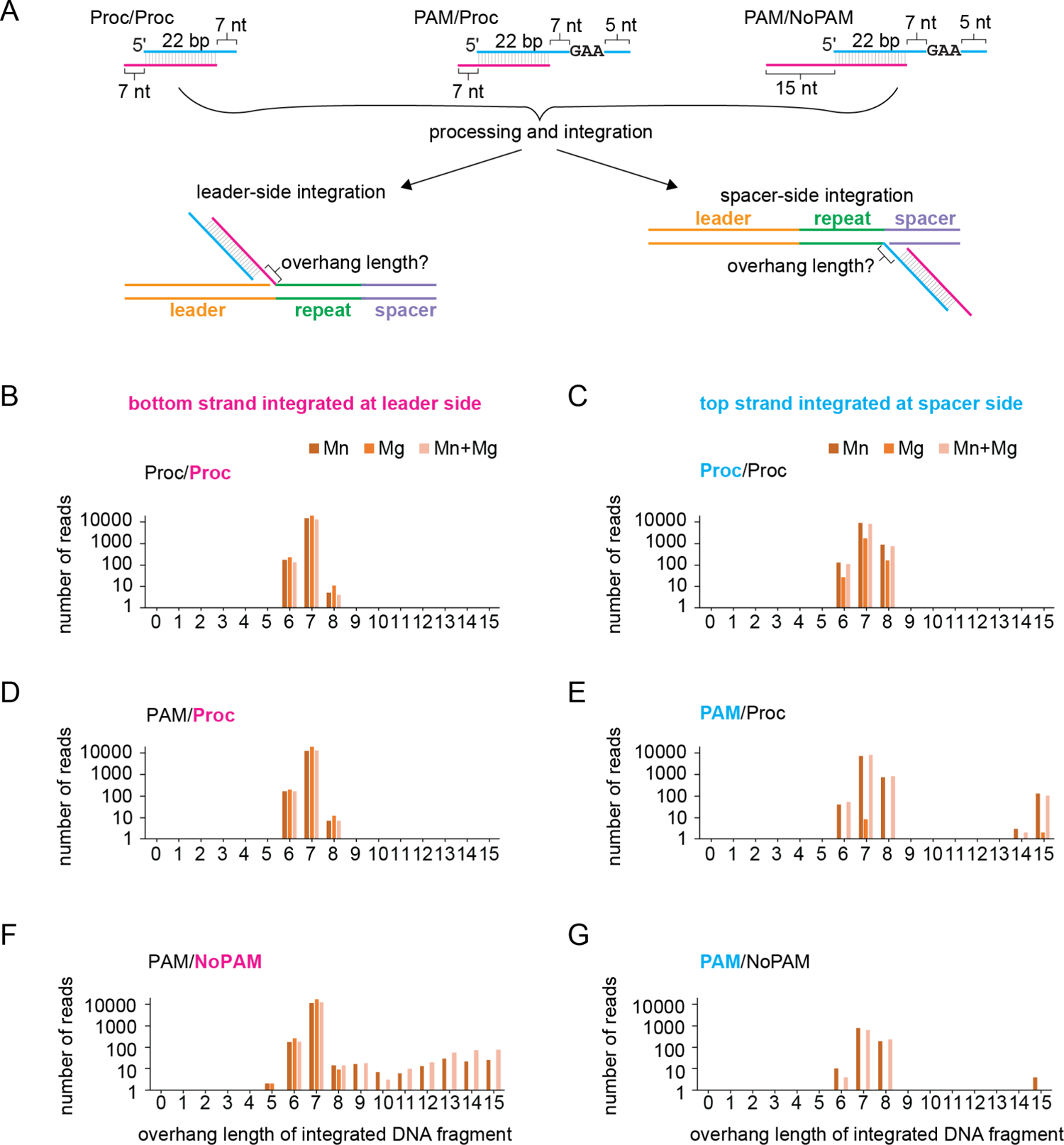
Cas1 cleavage products are integrated at the leader side. **A,** Substrate schematic of Proc/Proc, PAM/Proc and PAM/NoPAM substrates used in plasmid integration assays and integration products that were analyzed to determine overhang length following processing and integration. Top/PAM strand is shown in cyan and bottom/NoPAM strand is shown in magenta. **B-G,** Analysis of overhang length for processing/integration assays with substrates in (A) based on high-throughput sequencing of PCR Products shown in Fig. 4B. Number of reads are plotted against length of overhang for the bottom/non-PAM strand integrated at the leader side and the top/PAM strand integrated at the spacer side. Metal ion conditions were only 2 mM Mn^2+^, only 2 mM Mg^2+^, or both 1 mM MgCl_2_ and 0.5 mM MnCl. Note that the y-axis is plotted on a log scale.

For the pre-processed prespacer substrate (Proc/Proc) (**Fig. 5A**), overhang lengths ranged between 6-8 nt, with the actual overhang length of 7 nt observed in the vast majority of reads (**Fig. 5B-C**). The different overhang lengths may be due to sequencing errors, a small percentage of +/-1 oligonucleotide present in the prespacer substrate, or a small degree of Cas1-mediated prespacer processing, in the case of the 6 nt overhangs. The overhang length distribution was not dependent on the metal ions that were present during integration. Overhang length distribution for the pre-processed non-PAM strand in PAM-containing prespacer (PAM/Proc) was identical to the fully processed prespacer (Proc/Proc) (**Fig. 5B and 5D**).

When the PAM strand was integrated at the spacer side in the presence of Mn^2+^, most reads contained a 7 nt overhang, consistent with cleavage of the PAM strand just upstream of the GAA PAM sequence (**Fig. 5E and 5G**). The presence of some 6 and 8 nt overhangs could be due to sequencing errors, but also suggests a possibility of PAM “slipping” during processing by Cas4, generating variable length overhangs, consistent with in vivo analysis of Cas4-mediated spacer acquisition (43). Notably, we did not observe any 6 or 8 nt overhang integrated fragments in the absence of Mn^2+^, suggesting that these processing events were due to nucleolytic cleavage by the Cas4, and not the Cas1, domain.

Finally, for the prespacer substrate with an unprocessed non-PAM end (PAM/NoPAM), the vast majority non-PAM ends were processed to 7 nt in all metal conditions (**Fig. 5F**). This processing is consistent with cleavage by the Cas1 domain, which can use either Mn^2+^ or Mg^2+^ for its nuclease activity in a sequence-independent manner (**Fig. 1C and 2B**). The distribution of 6 to 8 nt overhang length for the non-PAM strand in the PAM/NoPAM substrate is similar to what we observe for the pre-processed strand in the other two substrates (**Fig. 5B and 5D**). There are a small percentage of reads for overhang length longer than 8 nt suggesting integration of some partially processed or unprocessed non-PAM ends (**Fig. 5F**). However, there are no reads with overhang lengths shorter than 5 nt, suggesting that the shorter cleavage products produced by Cas1 are not optimal for integration at the leader side (**Fig. 1C**), similar to our observation that short overhangs are not favorable for integration at the spacer side (**Fig. 3C, D**). Overall, these results reveal that the Cas1 domain can process non-PAM overhangs to an optimal length and catalyze their integration at the leader side, followed by PAM processing by the Cas4 domain and integration of the PAM end at the spacer side.

## Discussion

In this work, we demonstrate dual nuclease activities for a type I-G Cas4/1 fusion protein. The Cas4 domain recognizes PAM sequences and ensures that prespacers containing a PAM are captured and faithfully processed. The Cas1 domain processes the opposite end of the prespacer in a sequence non-specific manner, allowing generation of overhangs optimal for prespacer integration. Our plasmid integration assays provide evidence for directional integration of spacers by the Cas4/1 fusion. The non-PAM strand processed by the Cas1 domain is integrated at the leader side, while the PAM-strand cleavage products of the Cas4 domain are integrated at the spacer side. Structural and biochemical studies of the Cas4-Cas1-Cas2 and Cas4/1-2 complexes have shown that Cas4 protects the PAM end from exonucleases due to its strong affinity for PAM and blocks integration of the uncleaved PAM strand from being integrated by Cas1 (19, 22, 32). Our results are consistent with this model, as the PAM end was preferentially integrated at the spacer side following processing by the Cas4 domain.

Cas1 was identified as a nuclease in early biochemical studies of Cas1 proteins from type I-E *E. coli* and type I-F *P. aeruginosa* systems (40, 41), although a role for this nuclease activity remained unclear. These previous studies demonstrated cleavage of ssDNA, ssRNA, dsDNA and flapped and cruciform DNA substrates by Cas1 in a metal dependent and sequence non-specific manner. While *E. coli* Cas1 can use only Mg^2+^ (41), *P. aeruginosa* Cas1 is active in the presence of either Mn^2+^ or Mg^2+^ depending on the salt concentrations (40). Similarly, we observe that the Cas1 domain of type I-G *M. barkeri* Cas4/1 is active as a nuclease in the presence of either Mg^2+^ or Mn^2+^. We observe inhibition of the nuclease activity at high Mn^2+^ concentration. However, these inhibitory Mn^2+^ concentrations are higher than free Mn^2+^ concentrations in a bacterial cell suggesting that Cas1 could use either Mg^2+^ or Mn^2+^ within the cellular environment. Additionally, we observe a sequence independent mechanism for processing, allowing Cas1 to generate overhang lengths that are most favorable for integration regardless of the prespacer sequence. Notably, the Cas1 domain cleavage activity may not be generalizable to all type I-G Cas4/1 proteins, as a previous study of GsCas4/1 did not identify this activity (19).

Similar to previous studies (19, 22), we observed robust nuclease activity of the Cas4 domain of MbCas4/1 only in the presence of Mn^2+^ and at concentrations that were higher than physiological Mn^2+^ conditions. These results suggest that Cas4 may require additional activation to perform PAM cleavage in the presence of physiological metal conditions. It was previously shown that GsCas4/1 is activated for PAM cleavage following half-site integration, albeit in the presence of non-physiological Mn^2+^ concentration (19). However, we did not observe activation of MbCas4/1 PAM cleavage following half-site integration under any metal ion condition. Alternatively, an additional host factor may be required for Cas4 activation. Host factors like integration host factor (IHF) have been shown to aid in leader recognition by Cas1-Cas2 in type I-E and I-F systems (7, 14, 44–46). It is possible that interactions between similar CRISPR-bound host factors and Cas4-containing adaptation complexes assist with activation of Cas4 for PAM cleavage under physiological metal ion conditions.

Processing and directional integration of prespacers are key checkpoints of functional spacer acquisition. CRISPR systems use a variety of mechanisms to process non-PAM and PAM ends of prespacers and integrate these at the leader side and spacer side, respectively. In Cas4 containing systems, exonucleolytic processing of the non-PAM may enable rapid integration of at the leader side, followed by delayed PAM cleavage by Cas4 and spacer-side integration (19, 32). In Cas4-less systems, DnaQ and ExoT exonucleases, either standalone or fused to Cas2, have been shown to process both PAM and non-PAM overhangs in an asymmetric fashion (35–37, 39). In these systems, Cas1 contains an additional C-terminal domain that protects the PAM end from processing until leader side integration occurs (36). However, the mechanism of PAM cleavage in these systems remains unclear. The dual nuclease activities of type I-G MbCas4/1 fusion serves as an alternative mechanism for prespacer maturation. A nucleolytically active Cas1 domain that can process the non-PAM end of prespacers bypasses the need for host exonucleases to trim either end of the prespacer and makes the adaptation module self-sufficient for prespacer processing and integration.

### Experimental procedures

#### Cloning, protein expression and purification

Cas4/1 and Cas2 sequence from the type I-G system in *Methanosarcina barkeri str. Weismoor* were codon optimized and obtained as gBlocks from Integrated DNA Technologies (IDT). Cas4/1 was cloned into pET52b expression vector with a N-terminal 6x-His tag using Gibson assembly. Cas2 was cloned into pSV272 expression vector with an N-terminal 6x-His-MBP tag using Gibson assembly. Mutant constructs for Cas4/1, E101A and E375A were made using round-the-horn mutagenesis. sufABCDSE/pACYC construct was used for co-expression with Cas4/1 (32). All plasmids were verified using Sanger sequencing (Eurofins Genomics). All primers used for cloning various constructs have been listed in Table S1. Cas4/1 and Cas2 were expressed and purified using previously described procedures for Cas4 and Cas2 respectively (47).

#### Prespacer substrate preparation

All oligonucleotides used for cleavage and integration experiments were synthesized by IDT. Sequences of all substrates are shown in Table 2. All oligonucleotides were purified using 10% urea PAGE with 8M urea in 1x TBE followed by gel extraction and ethanol precipitation.

For radiolabeling, purified oligonucleotides were labeled on the 5′ end with [γ-^32^P]-ATP (PerkinElmer) and T4 polynucleotide kinase (NEB). Microspin G-25 columns (GE Healthcare) were used to remove the excess ATP from the labeling reaction. Duplex prespacer substrates were assembled by annealing a 5ʹ radiolabeled strand to an unlabeled complementary strand in a 1:2 ratio (25 nM radiolabeled strand and 50 nM unlabeled strand) in annealing buffer, 20 mM HEPES (pH 7.5), 100 mM KCl, 5% glycerol, 4 mM DTT. Hybridization reactions were heated at 95 °C for 5 min and cooled to room temperature.

#### Prespacer cleavage and half site integration assays

Cas4/1 and Cas2 were premixed on ice to a final concentration of 250 nM and substrate was added at a final concentration of 2-5 nM in reaction buffer, 20 mM HEPES (pH 7.5), 100 mM KCl, 5% glycerol, 4 mM DTT supplemented with MnCl_2_ or MgCl_2_ at concentrations indicated in the figures and figure legends. The reactions were incubated at 55 °C for 30 min. Reactions were quenched with 2X RNA loading dye (95% formamide, 0.01% SDS, 0.01% bromophenol blue, 0.01% xylene cyanol supplemented with 25 mM EDTA) and heated at 95 °C for 5 min. Samples were run on 10% urea-PAGE at 15 W for 1 to 1.5 hours. The gels were dried and imaged using phosphor screens on a Typhoon imager.

For single nucleotide resolution as in Figures 1C and 2B, reactions were ethanol precipitated with the addition of Glycogen Blue for easy detection of pellets after precipitation. Pellets were resuspended in 3 µL 2x RNA dye and 2 µL of the final resuspension was loaded on a 0.2 mm 8% urea-PAGE gel. The gel was run at 60 W for 1 to 2 hours, dried, and imaged using phosphor screens on a Typhoon Imager.

For quantification in Figure 3D, the intensity of bands was measured by densitometry using ImageJ (48). Percent integration was calculated by dividing the integration product band by the sum of both bands. The values from three replicates were averaged and error is reported as standard deviation between the replicates.

#### Plasmid integration assays

The leader and repeat sequences of the type I-G system were identified from the NCBI complete genome sequence CP009526.1 of *Methanosarcina barkeri str. Weismoor.* The sequence was used to design overlapping oligonucleotides, which were obtained from IDT. The oligo pairs were designed with sticky ends for BamHI and EcoRI. The oligo pairs were mixed and annealed in equal ratios, followed by ligation with double digested pUC19 at BamHI and EcoRI sites to obtain pCRISPR.

For the prespacers, purified oligonucleotides were annealed at a 1:1 ratio in annealing buffer with 20 mM HEPES (pH 7.5), 100 mM KCl, 5% glycerol at 95 °C for 5 min followed by cooling to room temperature.

Cas4/1 and Cas2 were premixed on ice to a final concentration of 250 nM in reaction buffer with 20 mM HEPES (pH 7.5), 100 mM KCl, 5% glycerol, 4 mM DTT supplemented with the divalent metal at concentrations indicated in the figures and figure legends. The prespacers were added at a final concentration of 500 nM and incubated on ice for 5 min. pCRISPR was added to the reaction at a final concentration of ∼15 ng/µl and the reactions were incubated at 55 °C for 1 h. Reactions were performed at a final volume of 100 µL. Reactions were cleaned up with Promega Wizard clean-up kit to remove salts, prespacers and proteins.

#### Sequencing sample preparation

2 µL of purified pCRISPR from the integration reactions was used for adaptor PCRs. 50 µl PCR reactions were performed with primers listed in Table S1 and Promega GoTaq enzyme. 34 cycles of 98 °C for 30 s, 58 °C for 30 s and 72 °C for 15 s were used for the PCR with initial denaturation at 98°C for 5 min and final extension at 72°C for 2 min. PCR products were analyzed on a 2% agarose gel pre-stained with SYBR Safe. 40 µL of the PCR was used for gel extraction. Bands corresponding to the expected size of 175 bp were excised and purified using Promega Wizard gel-extraction kit.

2.5 µL of purified adaptor PCR products were used for an additional PCR to add indices to samples for multiplexed MiSeq sample submission. 8 cycles of 98 °C for 30 s, 67 °C for 30 s and 72 °C for 10 s were performed for the PCR with initial denaturation at 98 °C for 30 s and final extension at 72 °C for 2 min. PCR products were analyzed on a 2% agarose gel pre-stained with SYBR Safe. Equal volumes of the final PCR reactions were mixed, and gel purified with the Promega Wizard gel-extraction kit. Sample quality was checked using Qubit and Agilent Bioanalyzer. Samples were sequenced using MiSeq with 2 x 75 cycles at the Iowa State University DNA Facility.

### Sequencing data analysis

R1 and R2 reads for demultiplexed fastq files were joined using ea-utils fastq-join (49). The sequences present in the resulting joined fastq files were analyzed using bash scripts. The total number of reads corresponding to an integration event in each joined fastq file was determined using one of the following commands:

1. grep -c GAGACGAGACAGTACACAACATGTG filename.fastq
2. grep -c CAAAGTGAATTCCGTAGCTAATAGACG filename.fastq These scripts search for the DNA sequence of the primer used to amplify (1) leader-side or (2) spacer-side integration products. These values were plotted in Fig. 4C as the total number of reads. To estimate the number of reads with a correct integration event at the leader side or the spacer side, we used grep to search for sequences corresponding to each end of the repeat followed by the expected overhang sequence.
3. grep -c CTTTCGGTTATGGAAACAAAAAA filename.fastq
4. grep -c TGTGGCAGAATTGAAGCAAAAAA filename.fastq

The script searches for (3) leader-side or (4) spacer-side integration products containing at least 6 nt overhangs. These values were plotted in Fig. 4C as the number of reads for correct integration events. The percent of correct integration events plotted in Fig. 4D were calculated by dividing the number of reads for correct integration events by the total number of reads.

To determine the length of overhangs for the L2 and S1 products, we used grep -c to search for the last five nt on either end of the repeat followed by all potential overhang sequences for a given prespacer sequence. The number of reads were plotted versus the overhang length for that sequence in Fig. 5B.

## Data availability

Any additional information, plasmids, reagents, and data are available from the lead contact at upon request.

## Supporting information

This article contains supporting information.

## Acknowledgements

We thank all members of the Sashital lab for helpful discussions and suggestions during this project. We thank the DNA facility, Office of Biotechnology at Iowa State University for MiSeq data collection.

## Author contributions

Y.D. and D.G.S. Conceptualization; Y.D. and D.G.S. Methodology; Y.D. and D.G.S. Investigation; Y.D. and D.G.S. Formal analysis; Y.D. and D.G.S. Visualization; D.G.S. Funding; D.G.S. Supervision; Y.D. Writing-original draft; Y.D. and D.G.S. Writing-review and editing.

## Funding and additional information

This work was supported by NIH R01 grant (GM115874) and NIH R35 grant (GM140876) to DGS.

