## Supplemental Information for "Cas4/1 dual nuclease activities enable prespacer maturation and directional integration in a type I-G CRISPR-Cas system"

**Supplementary Information**

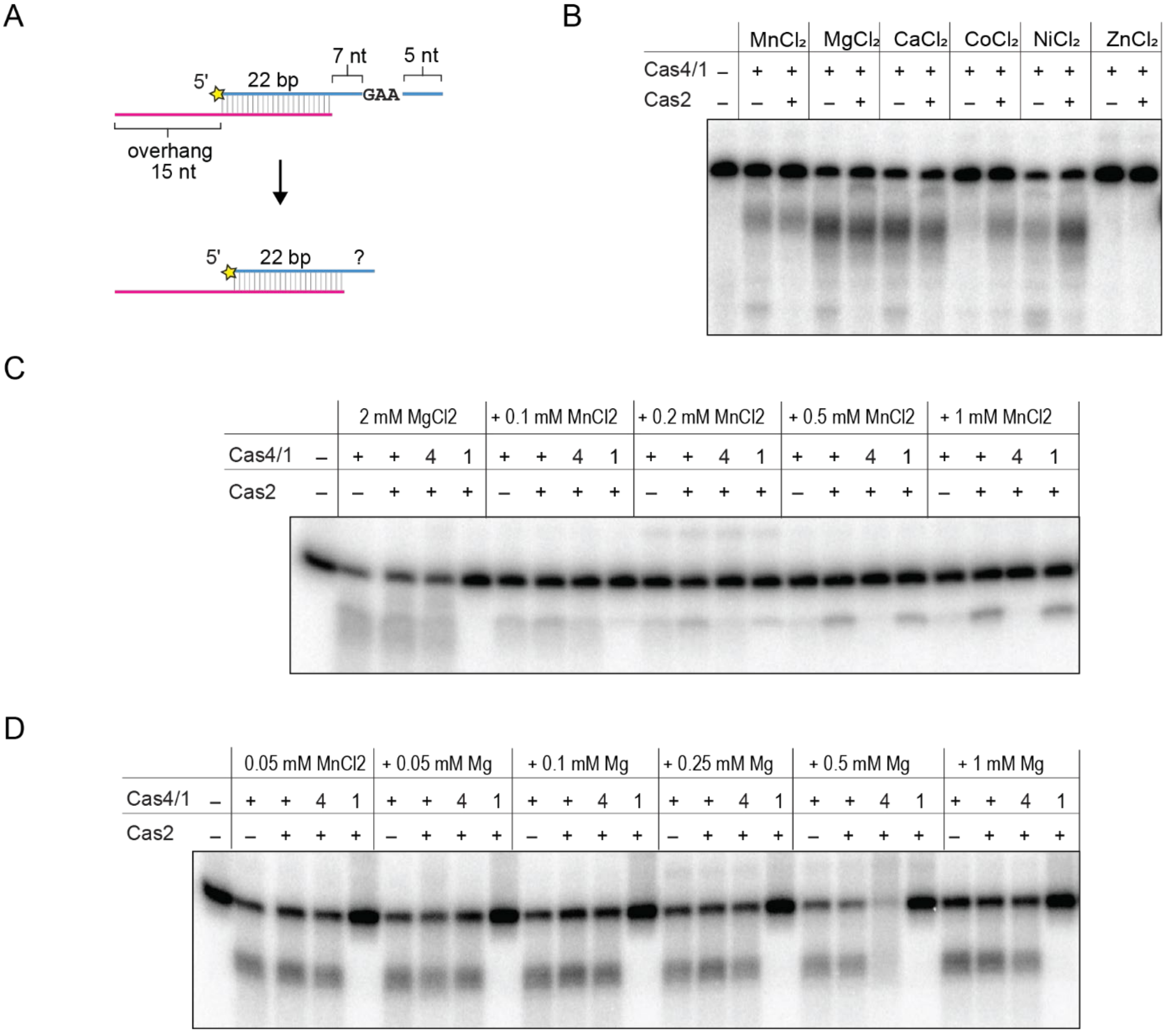


**Supplementary Figure 1. Cas4/1 nuclease activity in the presence of various metal ion cofactors**

**A,** Schematic of cleavage assay for a PAM/NoPAM substrate with the PAM strand radiolabeled**.** Radioactive label is indicated with a yellow star. **B,** Denaturing polyacrylamide gel showing cleavage assay with substrate shown in (A) in the presence of various metal ion cofactors. Metal ions were used at a final concentration of 2 mM. **C,** Cleavage assay for substrate shown in (A) with constant concentration of 2 mM $MgCl_2$ and increasing concentrations of $MnCl_2$ in the reaction. **D,** Cleavage assay for substrate shown in A with constant concentration of 0.05 mM $MnCl_2$ and increasing concentrations of $MgCl_2$ in the reaction.

**Supplementary Table 1. Primer used in this study**

| **Name** | **Sequence (5′ to 3′)** | **Description** |
|---|---|---|
| 1 | GTGGCGTACAAGAAGGGCTATGTACCTG | Forward for MbCas4/1 E101A |
| 2 | GGTACGGCTTTGCCGTTACTTTC | Reverse for MbCas4/1 E101A |
| 3 | ATTGCGGGAACTGCAGCCCAAATTTATTTTTCC | Forward for MbCas4/1 E375A |
| 4 | CCCCAATAACTGACCGATCTCC | Reverse for MbCas4/1 E375A |
| 5 | GATCCGAGACGAGACAGTACACAACATGTGAATGCCCATACTGTCCCTGGCTTCAATTCTGCCACAACCTTTCGGTTATGGAAACGGCGC | Oligo 1 for MbCRISPR assembly in pUC19 – complementary to oligo 4 |
| 6 | TAAGATCTTCACGTCTATTAGCTACGGAATTCACTTTGTTTTCGAGAGATCATTGAATTGAATTCTTTTCATGGATTATAAAACTAGCAT | Oligo 2 for MbCRISPR assembly in pUC19 – complementary to oligo 5 |
| 7 | ATTTATCTCAATTATAAAAGCTGAAGCTTCTCGAGAGCCTTCAGCAGTTTTTAGGGTTCATAAGCTCTCGAAAACG | Oligo 3 for MbCRISPR assembly in pUC19 – complementary to oligo 6 |
| 8 | AGGTTGTGGCAGAATTGAAGCCAGGGACAGTATGGGCATTCACATGTTGTGTACTGTCTCGTCTCG | Oligo 4 for MbCRISPR assembly in pUC19 |
| 9 | GAAAAGAATTCAATTCAATGATCTCTCGAAAACAAAGTGAATTCCGTAGCTAATAGACGTGAAGATCTTAGCGCCGTTTCCATAACCGAA | Oligo 5 for MbCRISPR assembly in pUC19 |
| 10 | AATTCGTTTTCGAGAGCTTATGAACCCTAAAAACTGCTGAAGGCTCTCGAGAAGCTTCAGCTTTTATAATTGAGATAAATATGCTAGTTTTATAATCCAT | Oligo 6 for MbCRISPR assembly in pUC19 |
| 11 | GTCTCGTGGGCTCGGAGATGTGTATAAGAGACAGCGTAGCTGAGGACCACCAGTAC | Prespacer top/PAM strand with Nextera adaptor – for L1 or S1 |
| 12 | GTCTCGTGGGCTCGGAGATGTGTATAAGAGACAGGTACTGGTGGTCCTCAGCTACG | Prespacer bottom/NoPAM strand with Nextera adaptor – for L2 or S2 |
| 13 | TCGTCGGCAGCGTCAGATGTGTATAAGAGACAGGAGACGAGACAGTACACAACATGTG | MbCRISPR backbone with Nextera adaptor – for L1 and L2 |
| 14 | TCGTCGGCAGCGTCAGATGTGTATAAGAGACAGCAAAGTGAATTCCGTAGCTAATAGACG | MbCRISPR leader with Nextera adaptor – for S1 and S2 |

**Supplementary Table 2. Oligonucleotides used in this study**

| Name | Sequence (5′ to 3′) | Description |
|---|---|---|
| 1 | CGTAGCTGAGGACCACCAGTACTTTTTTTGA ATTTTT | PAM strand for PAM/NoPAM or PAM/Proc prespacer (PAM 7 nt away from duplex) |
| 2 | GTACTGGTGGTCCTCAGCTACGTTTTTTTTTT TTTTT | NoPAM strand for PAM/NoPAM or prespacer |
| 3 | CGTAGCTGAGGACCACCAGTACTTTTTTT | Top strand for Proc/Proc prespacer |
| 4 | GTACTGGTGGTCCTCAGCTACGTTTTTTT | Bottom strand for Proc/Proc prespacer |
| 5 | CGTAGCTGAGGACCACCAGTACTTTGAATTT TTTTTT | PAM strand with PAM 3 nt away from duplex (Fig. 2) |
| 6 | CGTAGCTGAGGACCACCAGTACTTTTTGAAT TTTTTT | PAM strand with PAM 5 nt away from duplex (Fig. 2) |
| 7 | CGTAGCTGAGGACCACCAGTACTTTTTTTTTG AATTT | PAM strand with PAM 9 nt away from duplex (Fig. 2) |
| 8 | AGGACAACGTTACGGACGGCACAGCCTTTTT GAATT | PAM strand of prespacer for prespacer vs HSI cleavage assay (Fig. 3) |
| 9 | CCCTGTGCCGTCCGTAACGTTGTCGATTTTT | Proc strand of prespacer for prespacer vs HSI cleavage assay (Fig. 3) |
| 10 | CCCTGTGCCGTCCGTAACGTTGTCGATTTTTG TTTCCATAACCGAAAGGTTGTGGCAGAATTG AAGCGGCTTC | HSI substrate top strand with processed strand integrated at leader-side with repeat and spacer (Fig. 3) |
| 11 | GAAGCCGCTTCAATTCTGCCACAACCTTTCG GTTATGGAAACGGCGCTAAGATCTTTTGAT CTTAGCGCC | HSI substrate bottom strand with leader hairpin, repeat and spacer (Fig. 3) |
| 12 | AGGACAACGTTACGGACGGCACAGCCT | 3 nt overhang length strand for spacer side integration assay (Fig. 3) |
| 13 | AGGACAACGTTACGGACGGCACAGCCTT | 4 nt overhang length strand for spacer side integration assay (Fig. 3) |
| 15 | AGGACAACGTTACGGACGGCACAGCCTTT | 6 nt overhang length strand for spacer side integration assay (Fig. 3) |
| 15 | AGGACAACGTTACGGACGGCACAGCCTTTT | 6 nt overhang length strand for spacer side integration assay (Fig. 3) |
| 16 | AGGACAACGTTACGGACGGCACAGCCTTTTT | 7 nt overhang length strand for spacer side integration assay (Fig. 3) |
| 17 | AGGACAACGTTACGGACGGCACAGCCTTTTT | 8 nt overhang length strand for spacer side integration assay (Fig. 3) |
